# Biased inter-columnar communication and short-term plasticity in mouse barrel cortex

**DOI:** 10.1101/2025.11.04.686617

**Authors:** John M. Judge, Meyer B. Jackson

## Abstract

The barrel cortex (BC) processes complex direction-, frequency-, and phase-dependent input from whiskers to analyze objects in the immediate environment. Little is known about how BC microcircuits process this information and integrate input from multiple whiskers. To investigate these circuits we targeted a hybrid voltage sensor (hVOS) to Scnn1a excitatory neurons in cortical layer 4 (L4), and imaged population responses to electrical stimulation. BC in coronal and sagittal slices presented the laminar structure of the cortex with barrels aligned along stereotyped whisking directions. Voltage imaging tracked activity along an L4→L2/3→L4 relay during inter-barrel communication. AMPA receptor blockade demonstrated that this relay depends on excitatory synaptic transmission, and revealed intra- and inter-barrel feedforward inhibition. Communication between barrels was isotropic in response amplitude, half-width, and conduction velocity, but latency was longer for communication to caudal barrels. Furthermore, paired-pulse depression was weakest and recovery slowest for protraction-related directions, especially to caudally adjacent barrels. Such biases will preferentially enhance repetitive inputs in this direction. These results identify direction-dependent synaptic circuitry that shapes inter-barrel communication. Anisotropy in short-term plasticity aligns with whisker motion kinematics, suggesting that BC microcircuits are tuned to preserve temporal fidelity and selectively filter inputs according to whisking phase and direction.

---

The somatotopic map of mystacial whiskers in rodent primary somatosensory cortex provides a model for modular cortical processing (Egger et al., 2012; Stüttgen & Schwarz, 2018; Woolsey, 1970). Each whisker maps primarily to one barrel within the barrel cortex (BC), following the canonical thalamocortical path (Lübke et al., 2000; Viaene et al., 2011). How communication between barrels contributes to sensory processing remains poorly understood. Previous work has examined *in vivo* flow of sensory information (Armstrong-James et al., 1992; Zhang & Alloway, 2005) between layers and barrels, uncovering functional and morphological asymmetries between rows and arcs (Adesnik & Scanziani, 2010; Feldmeyer, 2012; Staiger & Petersen, 2021). However, corticothalamic feedback makes it difficult to isolate thalamocortical and cortical contributions to multi-whisker integration, particularly in L4 (Fox et al., 2003; Kwegyir-Afful et al., 2005; Lam & Sherman, 2009; Patel, 2019; Simons, 1978; Sun et al., 2006).

Whisking generates directional biases in sensory input that circuitry can process. Intracortical synapses may implement temporal filtering (Fortune & Rose, 2001) to operate on thalamocortical-transformed input at frequencies of 1-5 Hz for exploratory whisking to small-angle 15-25 Hz for foveal whisking (Adibi, 2019). Furthermore, these temporal preferences vary across the somatotopic map reflecting whisker specialization based on position (Hobbs et al., 2016; Woolsey, 1970). Studies show spatial or directional preferences to whisker deflections (Hubatz et al., 2020; Vilarchao et al., 2018), as well as phase-dependent processing during active whisking (Bermejo et al., 2002; Curtis & Kleinfeld, 2009; de Kock & Sakmann, 2009; Kleinfeld et al., 2016; Leiser & Moxon, 2007). However, little is known about the inter-barrel microcircuits mediating these anisotropies.

Experiments in brain slices reveal intra-cortical communication in isolation from sensory input, and have shown that electrical stimulation evokes activity that propagates from L4 to supragranular layers before spreading to other barrels (Hill & Greenfield, 2014; Laaris & Keller, 2002; Petersen & Sakmann, 2001; Sato et al., 2008). Here we targeted a hybrid voltage sensor (hVOS) to L4 excitatory neurons (spiny stellate and pyramidal) in mouse BC, and prepared cortical slices to image voltage in the neurons that receive the primary thalamic input. Excitatory synapse blockade isolated barrels and revealed that communication between barrels depends on synaptic transmission in L2/3 and L4. We investigated response characteristics, velocities, and synaptic depression in BC microcircuits, and evaluated specific inter-barrel connections to gain insight into isotropic and anisotropic aspects of inter-barrel communication.

## Materials and Methods

### Animals

Ai35-hVOS1.5 (C57BL/6-Gt(ROSA) 26Sortm1 (CAG-hVOS1.5)Mbja/J; JAX #031102) Cre reporter mice were bred with Scnn1a-tg3-Cre driver mice (B6;C3-Tg(Scnn1a-cre)3Aibs/J; JAX # 009613) to create Scnn1a::hVOS mice with hVOS probe expression (Bayguinov et al., 2017; Chanda et al., 2005) in L4 excitatory neurons (Madisen et al., 2009; Scala et al., 2019). All animal procedures were approved by the Animal Care and Use Committee of the University of Wisconsin-Madison School of Medicine and Public Health (protocol: M005952).

### Slice preparation

Male and female mice were deeply anesthetized with isoflurane and sacrificed via cervical dislocation. Dissected brains were placed into ice-cold cutting solution (in mM: 10 glucose, 125 NaCl, 4 KCl, 1.25 NaH_2_PO_4_, 26 NaHCO_3_, 6 MgSO_4_, 1 CaCl_2_) bubbled with 95% O_2_/5% CO_2_. We cut ∼6 coronal or ∼4 sagittal 200-μm slices per brain hemisphere, with a Leica VT1200S vibratome and placed them into 95% O_2_/5% CO_2_-bubbled artificial cerebrospinal fluid (ACSF, same as cutting solution except 1.3 mM MgSO_4_, 2.5 mM CaCl_2_, 4 μM DPA) for at least 1 h (Scheuer et al., 2023).

### Voltage imaging and electrical stimulation

Slices continuously perfused with ACSF at room temperature were viewed using an Olympus BX51 microscope and XLUMPlanFl 20× objective (N.A. 1.0). A triggered pulse train generator (Prizmatix, Holon, Israel) and stimulus isolator (World Precision Instruments, Sarasota, FL) generated stimulus pulses (50-250 μA, 200 μs) applied via fire-polished, ACSF-filled KG-33 glass microelectrodes (King Precision Glass, Claremont, CA) with tip diameters ∼6–8 μm. Displayed traces of fluorescence versus time were averages of five trials, at 15-second intervals.

### Voltage imaging: cameras and software

Slices were illuminated by an LED (435-nm peak emission; Prizmatix, Holon, Israel). Gradient contrast images were acquired with a high-resolution Kiralux CMOS camera (Thorlabs, Newton, NJ). For voltage imaging we used a da Vinci 2K CMOS camera (2 kHz, 512×160 pixels binned/cropped to 80×80; RedShirt Imaging, Decatur, GA). A movable mirror and dual-port adapter switched between the cameras. Acquisition was controlled by TurboSM (RedShirt Imaging/SciMeasure) and a C++ application for data processing, PhotoZ (Chang, 2006). A Python application (OrchestraZ) automated the workflow.

### Analysis and Statistics

We analyzed data from 11 male (21 slices, 48 barrels) and 8 female (16 slices, 39 barrels) mice aged 4-13 weeks aged 7.8 ± 1.7 weeks (2-3 barrels per slice, 1-4 slices per animal). Age and sex analyses are included in Supplementary Results. Data processing and statistical significance tests are detailed in Supplementary Methods.

## Results

We characterized barrel organization in coronal and sagittal slices according to Scnn1a labeling, and evaluated the alignment between barrels and whiskers. We used voltage imaging to follow the spread of excitation within and between barrels, evaluating the contributions of excitatory synaptic transmission to inter-barrel communication along an L4→L2/3→L4 relay. We then determined response parameters of inter-barrel communication in different directions.

Finally, we assessed short-term plasticity between barrels along behaviorally relevant axes.

### Barrel visualization and alignment

The BC columnar organization is visible in low magnification fluorescence images of somatosensory cortex slices from Scnn1a::hVOS mice (Fig. 1). Labeling is concentrated in L4 barrel hollows. We used the Brain Globe atlas of mouse BC (Claudi et al., 2020) to evaluate directionality of inter-barrel communication during voltage imaging. Coronal slices preserved dorsal-ventral connections (Fig. 1A-C) and sagittal slices preserved rostral-caudal connections (Fig. 1D-F). However, these slices cannot strictly preserve rows and arcs due to the curvature of the barrel field grid, and its rotation from the whisker grid. This is consistent with typical medial-lateral angles used to cut tangential BC slices (Sumser et al., 2017). Accounting for this alignment, we categorized our slices based on aspect and orientation.

**Fig. 1:**
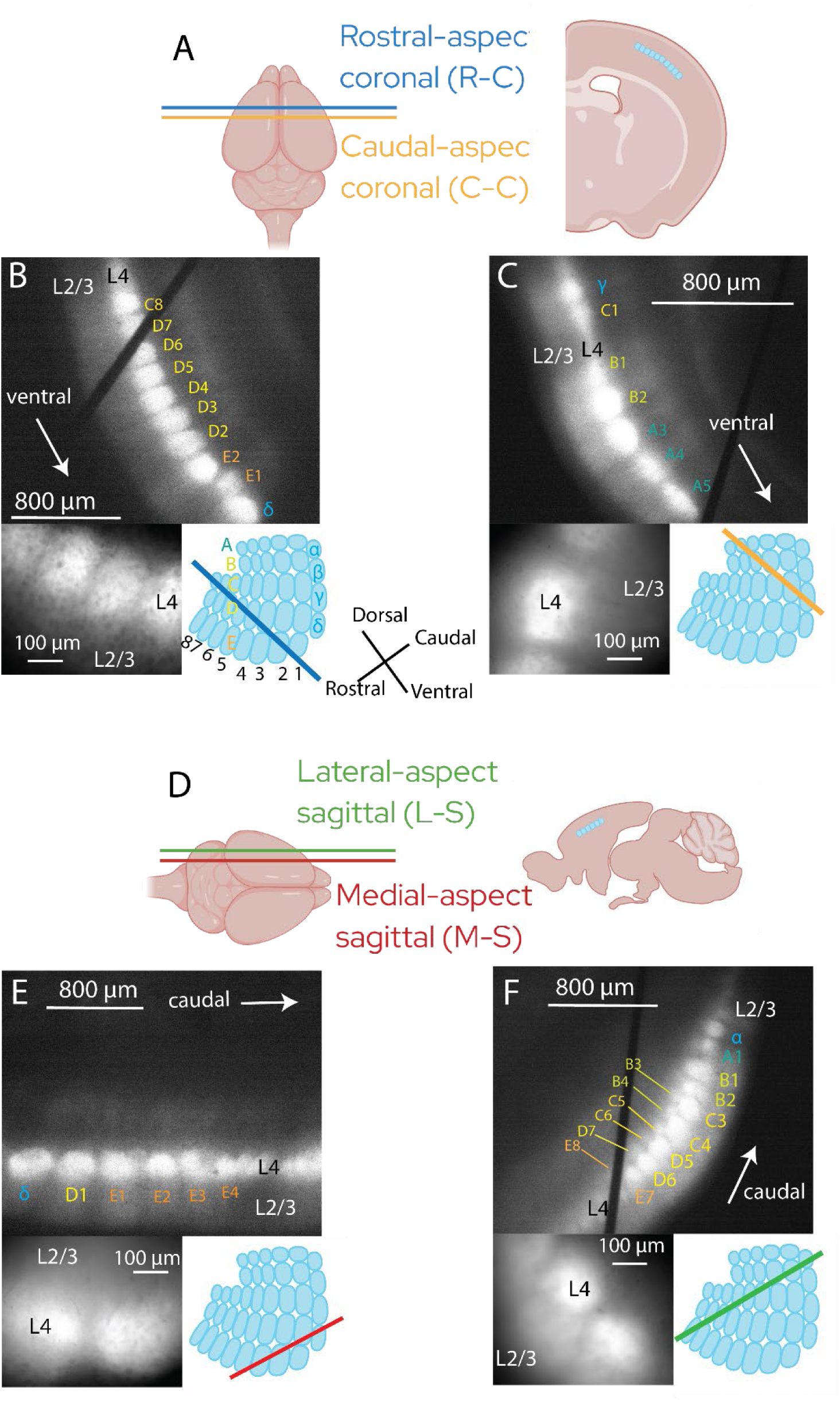
Anatomical orientation and slicing strategy. **A.** Left: dorsal view of mouse brain in the coronal plane: rostral-aspect coronal (R-C, blue) and caudal-aspect coronal (C-C, orange) sections. Right: Coronal slice with barrels highlighted as blue (created with Biorender). **B-C**. Top: low-magnification fluorescence and gradient contrast images superimposed from R-C (B) and C-C (C) slices, with annotations for barrel, barrel relationship, and orientation. Lower left: higher magnification of the corresponding slice. Lower right: diagram of tangential section of barrel cortex; blue and orange lines indicate approximate location of respective slicing planes for R-C and C-C in the barrel field. **D.** Left: Dorsal view of mouse brain showing slicing planes: lateral-aspect sagittal (L-S, green) and medial-aspect sagittal (M-S, red). Right: illustration of barrels in a sagittal slice (created with Biorender). **E.** M-S slices; **F**. L-S slices, paralleling panels and annotations in B and C. Barrel row and arc assignments are given to within one row or arc; this uncertainty was acceptable because determining only aspect and plane classification is needed. Note that the black fibers visible in the low magnification images are threads of the harps used to anchor slices.

Figs. 1B-C and 1E-F show barrel relationships annotated (top), accompanied by a higher magnification image (bottom left) and a tangential view of BC with slice plane drawn (bottom right). We divided slices into four groups: rostral-aspect coronal (R-C slice, Fig. 1A, 1B), caudal-aspect coronal (C-C slice, Fig. 1A, 1C), lateral-aspect sagittal (L-S slice, Fig. 1D, 1E), and medial-aspect sagittal (M-S slice, Fig. 1D, 1F). L-S slices were excluded from the present study because of their ambiguous organization (see Supplementary Results). The other sections align well to specific stereotyped whisking patterns. R-C and C-C slices track exploratory whisking, with protraction angles to the rostral-dorsal direction. M-S slices track foveal whisking, with protraction angles toward the rostral-ventral direction (Knutsen et al., 2005; Petersen et al., 2020). Basic barrel dimensions further distinguish the R-C, C-C, and M-S slices. Septal width, barrel width, and barrel height varied between these slice categories (see Fig. S1 in Supplementary Results), providing further correspondence with inter-barrel relations.

### Stimulation induced depolarization of multiple barrels

In both coronal and sagittal slices, electrical stimulation in L4 elicited fluorescence changes in both the *home* (stimulated) barrel, and the two *neighboring* barrels (Fig. 2). A coronal slice was stimulated with currents ranging from 50 to 250 μA, and maps with color-coded response amplitude (peak ΔF/F) illustrate the spatial spread of depolarization (Fig. 2A-E). To track the evolution of responses, we plotted mean fluorescence within selected ROIs versus time (Fig. 2A-E, traces below each map). Since hVOS is a negative-going voltage indicator (Chanda et al., 2005), these downward-going signals report depolarization of probe-expressing neurons. Responses in neighboring barrels have smaller amplitudes and slower time courses compared to the home barrel. The response amplitudes in the home (blue traces) and neighbor barrels (orange and pink traces) increased with stimulation current and saturated with strong stimulation.

**Fig. 2:**
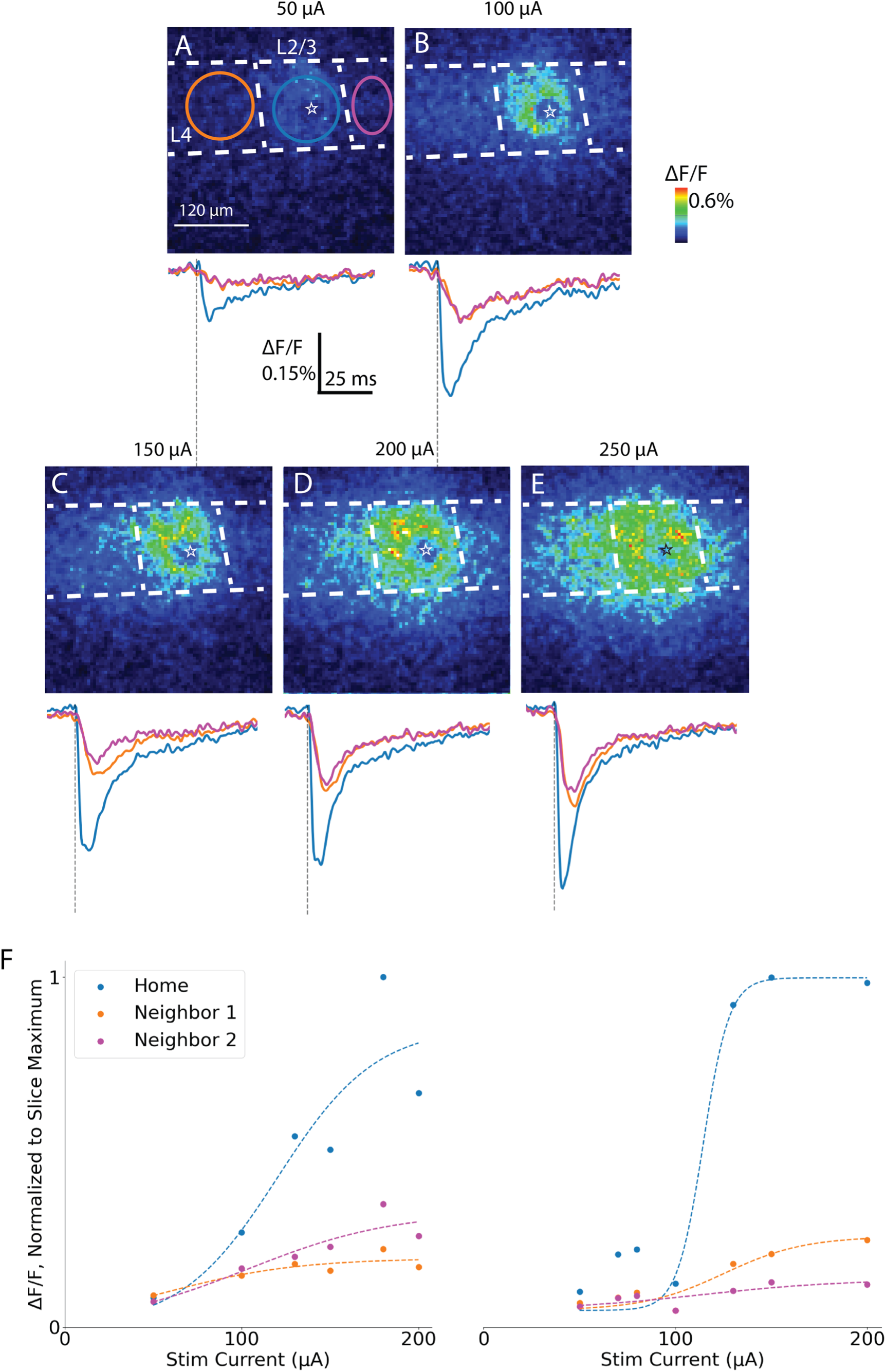
Stimulation-induced voltage changes in barrel cortex slices. **A.** Map of peak ΔF/F in response to 50 μA stimulation current in coronal slice with L4 stimulation site marked with a star. ROIs within home and neighbor barrels are delineated with colored boundaries. **B-E.** Maps of peak ΔF/F in response to increasing stimulation currents (color scale denotes response amplitude). Traces below each panel show fluorescence versus time for each ROI (colors of traces correspond with ROI outlines). Spatial scale bar in **A** applies to all images. **F**. Plots of peak ΔF/F versus stimulation current from two different experiments for home and neighbor barrels. Plots were fitted with sigmoid curves (see text, dotted lines) for the home and neighbor barrel plots.

We plotted peak ΔF/F versus current and fitted the curves to a sigmoidal function of the form 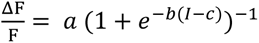 with stimulation current 𝐼𝐼 (Fig. 2F: two examples). This fit the data well for 7 of 8 home barrels (home R² = 0.93 ± 0.03) and 12 of 14 neighbor barrels (R² = 0.88 ± 0.03). We determined mean 70%-saturation stimulation (home: 129 ± 11 μA; neighbor: 178 ± 26 μA) and used this current determined for each slice (ranging from 80-200 μA) to activate inter-barrel circuitry.

### Barrel responses to stimulation of different layers

Stimulation in L4 produced less spread than stimulation in L2/3 (Figs. 3A-B versus Figs. 3D-E). Responses in neighboring barrels had smaller amplitudes for L4 (Fig. 3C) versus L2/3 (Fig. 3F) stimulation. Furthermore, neighboring and home barrel responses had similar amplitudes with L2/3 stimulation, but neighboring barrels had smaller response amplitudes with L4 stimulation. The ratio of neighbor peak ΔF/F to home peak ΔF/F was significantly less than one with L4 stimulation, 0.52 ± 0.13 (Fig. 3G, 1-sample t-test on neighbor/home ΔF/F ratio: T = 3.7, p = 0.003, n = 9 slices) but indistinguishable from one with L2/3 stimulation, 0.97 ± 0.28 (T = 0.09, p = 0.5, n = 6). Together, these data suggest that L4 responded more strongly to L2/3 stimulation during inter-barrel communication.

**Fig. 3:**
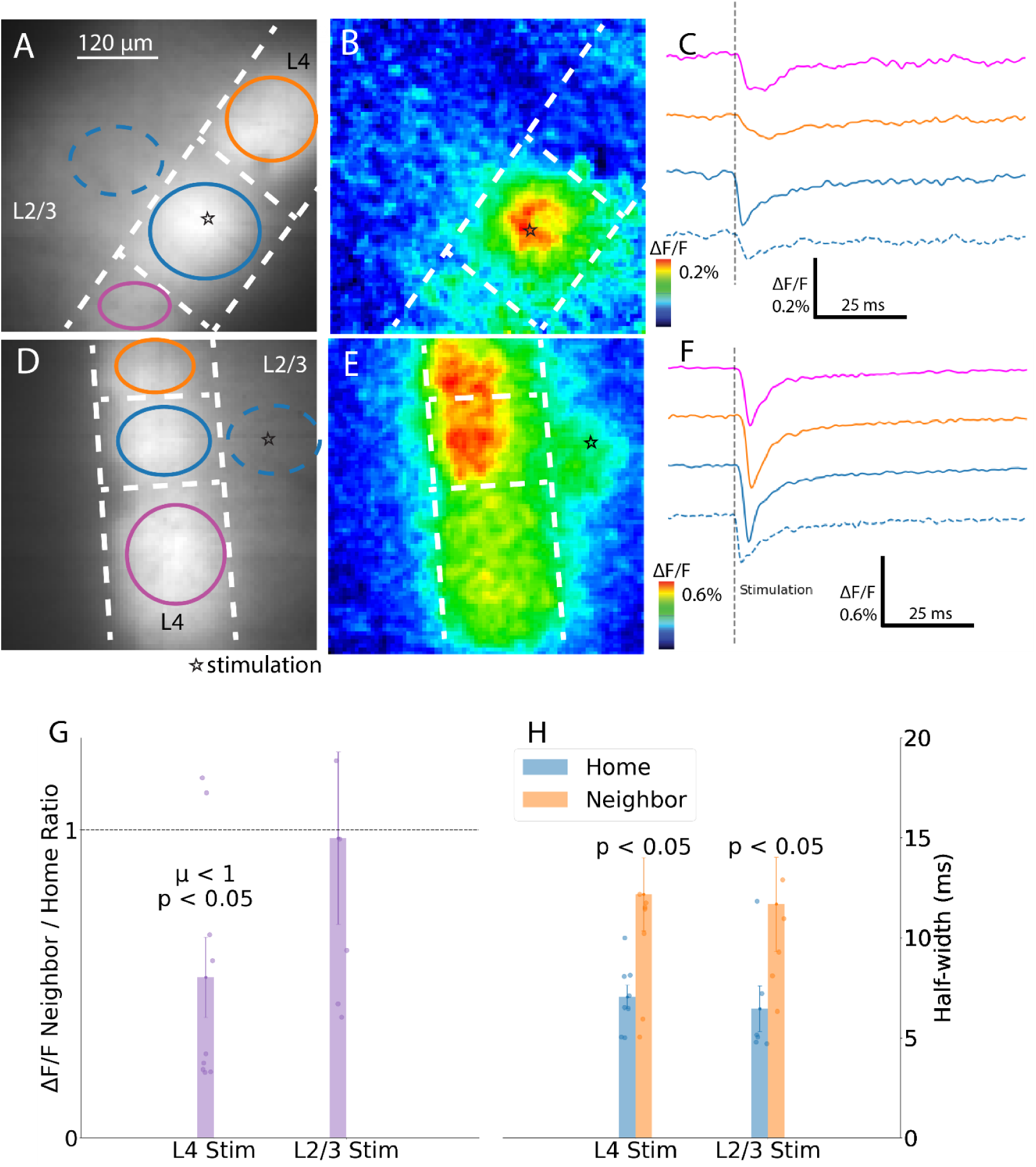
Response patterns for stimulation in L4 and L2/3. **A-C**. L4 stimulation evokes strong responses in home barrels and weaker responses in neighbor barrels. **A**. Image of coronal slice with star marking the stimulation site. ROIs drawn within home and neighbor barrels. **B**. Map of peak ΔF/F (color scale denotes response amplitude; each map is scaled to its maximum). **C**. Traces of ROI-averaged fluorescence intensity with corresponding color and line style as in A. **D-F.** L2/3 stimulation elicits nearly equal responses across barrels. D-F parallel A-C. **G.** Neighbor/home ratios of peak ΔF/F are significantly lower for L4 than L2/3 stimulation (n = 9 and 6 barrels). **H**. Neighbor barrel half-widths were significantly greater than home barrel for both stimulation sites. Bars: mean ± SEM with points for individual slices.

L4 and L2/3 stimulation evoked neighbor barrel responses with significantly greater half-widths by almost a factor of two (Fig. 3H, Welch’s t-test, L4 stimulation, home: 8.8 ± 0.7, neighbor: 15.2 ± 2.3 ms, T = 2.7, p = 0.01; L2/3 stimulation, home: 8.1 ± 1.4, neighbor: 14.6 ± 2.9 ms, T = 2.0, p = 0.04). We also saw small fluorescence changes in L2/3 (dotted blue for the home barrel in Fig. 3E). Because the probe is expressed in neurons with somata located in L4 (Madisen et al., 2009), we interpreted these responses in L2/3 as depolarization of axons and dendrites of our targeted Scnn1a neuronal subpopulation.

### AMPA-receptor blockade identifies postsynaptic responses

The spread of excitation illustrated in Fig. 3 reflects both direct depolarization and orthodromic responses. Because orthodromic responses depend on excitatory synapses we assessed their contribution by blocking AMPA receptors with the antagonist NBQX (10 μM). Fig. 4A illustrates home and neighboring barrel responses to L4 stimulation. NBQX nearly eliminated the depolarization of the neighbor barrel (Fig. 4B, left), reducing the peak amplitude by more than 5-fold (Fig. 4B, right). This strong blockade demonstrated the dependence of these depolarizations on AMPA receptor-mediated excitatory synapses. By contrast, NBQX increased response amplitudes within the home barrel (Fig. 4B), suggesting these responses were largely direct. Increases in response amplitude indicated that local inhibitory interneurons, activated by excitatory synapses, limited the responses seen in ACSF alone. NBQX had almost no inhibitory action on either home or neighbor barrel response amplitudes when L2/3 was stimulated, indicating that antidromic action potentials dominate these responses (Fig. 4C-D).

**Fig. 4:**
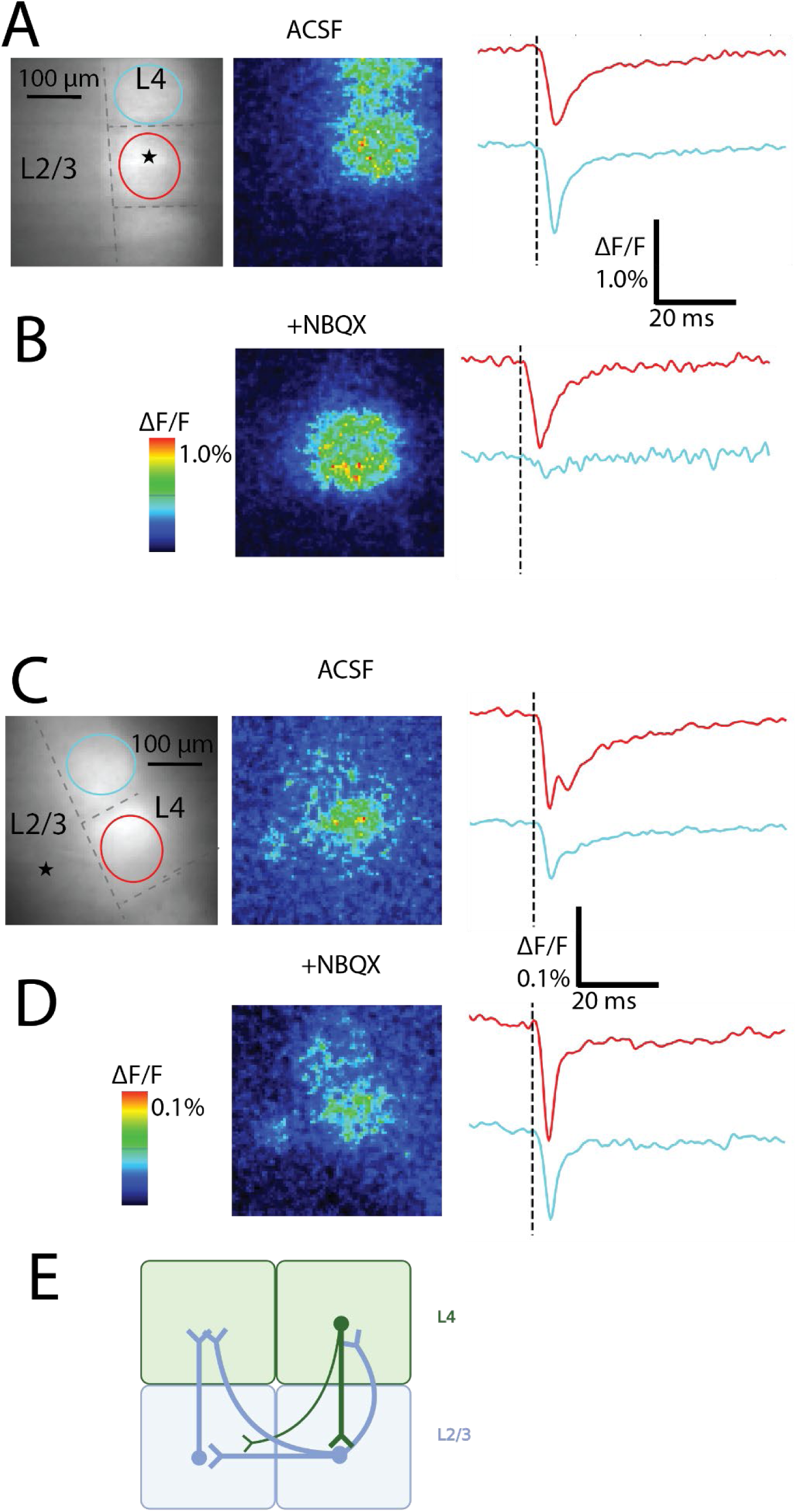
AMPA receptor blockade removes synaptic response components. **A.** L4 stimulation elicits responses in home and neighbor barrels. Left: resting fluorescence image of coronal slice with L4 stimulation site and ROIs annotated. Center: Map of peak ΔF/F. Color scale denotes response amplitude. Right: Traces of fluorescence versus time for each ROI of the corresponding color in the left panel. Vertical black dashed line: stimulation time. **B.** NBQX strongly inhibited neighbor barrel responses, and slightly increased home barrel responses. Color scale applies to maps in A and B. Center/right panels parallel those in A. **C-D.** With L2/3 stimulation, NBQX had little effect on amplitude but reduced half-width. C-D parallel A-B. **E.** Circuit diagram: L4→L2/3→L4 mediates inter-barrel spread from L4 stimulation via monosynaptic or disynaptic pathways or both. Processes of Scnn1a neurons in L2/3 are activated antidromically by L2/3 stimulation. Thickness of axons in the diagram reflects the density of these connections in reconstruction studies (Staiger & Petersen, 2021) (Created with Biorender).

We interpreted these results with a minimal circuit (Fig. 4E) in which L4 stimulation first activates excitatory neurons in L4. These neurons then excite non-Scnn1a excitatory neurons with cell bodies in L2/3, which in turn excite Scnn1a L4 excitatory neurons in the neighboring barrel. This circuit illustrates how intracortical multi-barrel activation uses L2/3 as a relay.

Responses of neurons residing in L2/3 to stimulation of L4 depend on excitatory synaptic transmission at L4→L2/3 synapses or antidromic activation of the axons of L2/3 neurons extending into L4. L2/3 neurons thus activated then excite L2/3 and L4 neurons in the neighboring barrels. A few axons from L4 neurons are reported to pass through L2/3, cross into the adjacent barrel, then curve back up to L4. However, inter-barrel axons originating in L2/3 outnumber these direct L4→L4 inter-barrel axons (Staiger & Petersen, 2021), consistent with the strong L4 inter-barrel signaling blockade by NBQX observed here. Meanwhile, L2/3 stimulation directly activates the extensive L2/3 inter-barrel arborization, which conducts action potentials antidromically to L4 neurons in multiple barrels.

To examine spatial variations in the action of NBQX at a finer scale, we constructed difference maps comparing responses before and after NBQX, with two examples each for stimulation in L4 (Fig. 5A) and L2/3 (Fig. 5B). Difference maps (denoted with “Δ”) display the change in peak ΔF/F from ACSF to NBQX for each pixel (white: no change; red: increase; blue: decrease). These difference maps show that NBQX reduced responses to L4 stimulation (Fig. 5A, darkest blue under “Δ”) outside the home barrel, while home barrel responses were unchanged or increased. Difference maps for L2/3 stimulation showed that NBQX broadly increased responses in L4 of both barrels (Fig. 5B). We plotted differences for pixels in the neighboring barrel as histograms (to the right of difference maps). Neighbor barrel distributions had means significantly less than zero for L4 stimulation (one-tailed 1-sample t-test, L4, top: T = −8.7, p < 0.001, middle: T = −38, p < 0.001, bottom: T = −33, p < 0.001) but not for L2/3 stimulation (top: T = 15, p = 1.0, middle: T = 9.4, p = 1.0, bottom: T = 6.1, p = 1.0). Using a slice-specific ΔF/F threshold (70^th^ percentile of recording pixels in ACSF) we calculated the number of responding pixels (36 μm^2^ per pixel). The area responding to L2/3 stimulation was not significantly different between NBQX and control (Welch’s t-test, T = 0.15, p = 0.88, 6 slices), while with L/4 stimulation NBQX significantly decreased response area (Fig. 5C, T = 2.8, p = 0.018). NBQX also significantly decreased peak ΔF/F of neighbor barrels for L4 stimulation (T = 3.1, p = 0.01), but the decrease was not significant for L2/3 stimulation (Fig. 5D, T = 0.08, p = 0.94).

**Fig. 5:**
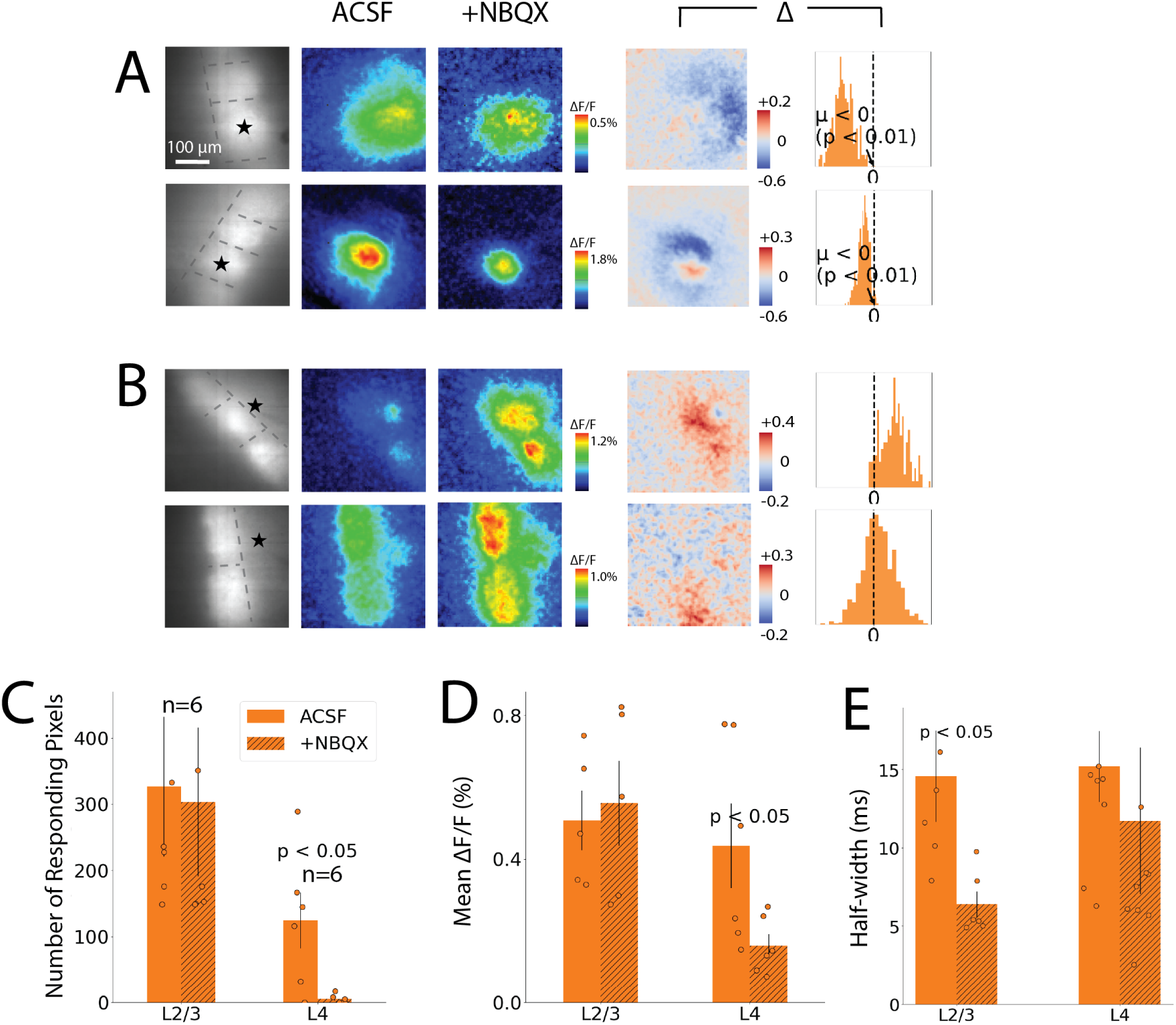
Spatial patterns of NBQX action. **A-B**. Each row corresponds to one slice before and after adding NBQX, with L4 stimulation (A) and L2/3 stimulation (B). Left-most column: resting fluorescence images with stimulation site (star) and layer/barrel boundaries (dashed lines). ACSF column: Maps of peak ΔF/F in ACSF. +NBQX column: Maps of peak ΔF/F after NBQX addition. Color scale matches maps in ACSF column (difference scale for each row). Δ column, left: Difference maps (Δ = NBQX − ACSF) show suppression in neighbor barrels for L4 but not L2/3 stimulation. White pixels represent zero change; red pixels represent responses increased by NBQX, and blue pixels represent responses reduced by NBQX. Δ column, right: histograms of pixelwise differences for neighboring barrel only. Means were significantly less than zero in both examples for L4 stimulation (p < 0.01; one-sample t-test; A top: −0.20 ± 0.03; A bottom: −0.73 ± 0.02), but not for L2/3 stimulation (B top: 0.12 ± 0.06; B bottom: 0.03 ± 0.01). **C.** Responding pixels were defined as pixels in the neighboring barrel with peak ΔF/F above a slice-specific threshold. Response area (based on number of pixels) was decreased by NBQX for L4 but not for L2/3 stimulation. **D**. Mean peak ΔF/F was reduced by NBQX in neighbor barrels for L4 but not for L2/3 stimulation. **E.** Half-width before and after NBQX for neighbor barrels. Bars show mean ± SEM, with points for individual slices.

Responses in the absence of NBQX appeared biphasic (Fig. 4C). NBQX removed the second component (Fig. 4D), suggesting that synaptic delays of several milliseconds separated synaptic from direct or antidromic responses. Response half-widths reflect this separation. In neighboring barrels NBQX reduced half-widths more than two-fold for L2/3 stimulation from 14.6 ± 2.9 to 6.4 ± 0.8 ms (Fig. 5E; Welch’s t-test, L2/3 neighbor barrel: T = 2.7, p = 0.02). By contrast, for L4 stimulation the reduction (15.2 ± 2.3 to 11.7 ± 4.7 ms) was not significant (Fig. 5E; L4 neighbor barrel: T = 0.7, p = 0.5).

### Propagation through layers and barrels

We further tested the hypothesis that inter-barrel L4→L4 communication depends on conduction through L2/3 by following the propagation of responses along this route. For the slice shown in Fig. 6A, with a peak response map in Fig. 6B, the traces in Fig. 6C illustrate this sequence. Responses in the L4 home barrel are followed first by responses in L2/3 of the home barrel, and then by responses in L4 of the neighboring barrel. Stimulating L2/3 (Fig. 6D) produced the response heatmap in Fig. 6E, and the traces in Fig. 6F illustrate nearly simultaneous activation of L4 in the home barrel along with L4 and L2/3 in the neighboring barrel.

**Fig. 6:**
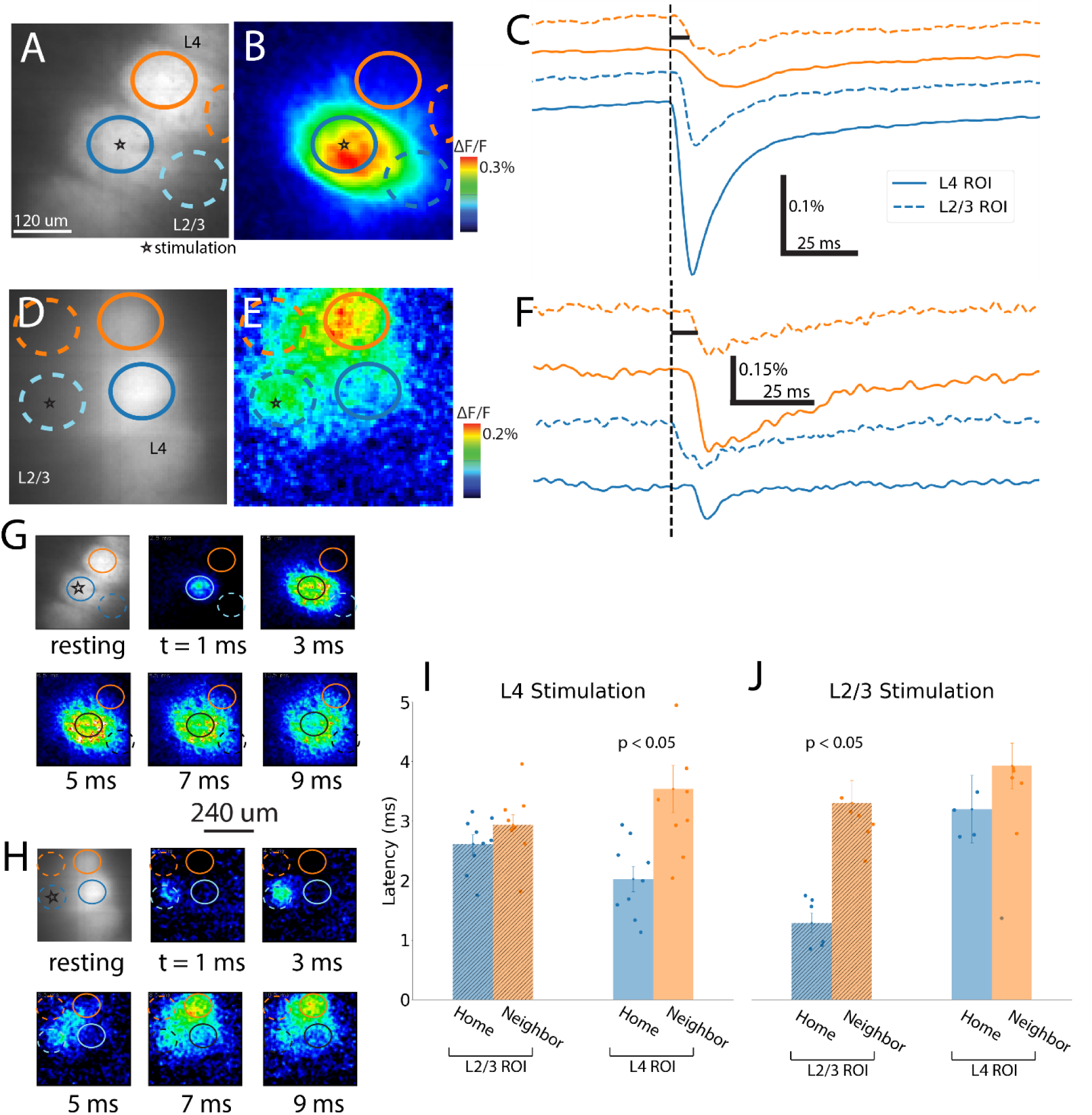
Spatiotemporal spread of excitation across barrels and layers. **A** and **D**. Fluorescence images showing ROIs and stimulation sites. **B** and **E.** Heat maps of peak responses (color scale denotes amplitude). **C** and **F.** Traces in the home (blue) and neighbor (orange) barrels of L4 (solid) and L2/3 (dotted). ROI outlines in maps correspond to traces with corresponding colors. Dashed black line marks stimulation time. With L4 stimulation, the response in the L4 home barrel precedes that in L2/3 and in neighbor barrels. **D-F.** Stimulation in L2/3 elicits near-simultaneous activation of three nearby ROIs following responses in the stimulated L2/3 ROI. **G-H.** Upper-left: fluorescence images corresponding to A and D, with successive panels providing a time-lapse sequence of ΔF/F maps at 1 ms intervals following stimulation in L4 (G) or L2/3 (H) to illustrate the spread of excitation over a 9 ms window. **I-J.** Latency to half-maximum ΔF/F for each ROI averaged over multiple slices for L4 stimulation (n = 10) and L2/3 stimulation (n = 7). Means are plotted ± SEM with points for individual slices. L4 stimulation leads to sequential activation of L4 in home followed by neighbor barrels, while L2/3 stimulation leads to more synchronous activation across regions.

Amplitude maps at regular time intervals chart the spread of excitation evoked by stimulation of L4 (Fig. 6G) and L2/3 (Fig. 6H). Note that these maps are snapshots in time, as opposed to peak response maps displayed in the preceding figures. The time from stimulation to half-maximum on the rising phase, or *latency,* was 1.9 ms for L4 home barrel responses to L4 stimulation, followed by the L2/3 home barrel at 3.0 ms and the L2/3 neighbor barrel at 3.9 ms (Fig. 6G). L4 of the neighbor barrel responded with the longest latency (6.2 ms), reflecting synaptic activation of L2/3 excitatory neurons, conduction through their axons, and synaptic activation in L4. With L2/3 stimulation, local responses in L2/3 appear first (1.5 ms), followed within 3-4 ms by essentially simultaneous activation of the other three regions, starting in the 5-ms snapshot (Fig. 6H). Multimedia 1 presents another example of spread of responses to L4 stimulation, illustrating the L4→L2/3→L4 conduction route. Multimedia 2 presents another L2/3 stimulation example, illustrating the simultaneous L2/3→L4 antidromic conduction and biphasic responses (Fig. 4D, traces). Mean latencies also support these sequences for L4 (Fig. 6I, n = 10) and L2/3 stimulation (Fig. 6J, n = 7). Latencies differed significantly among the four ROIs (ANOVA; Fig. 6I: F = 6.3, p = 0.0017; Fig. 6J: F = 7.9, p = 0.0010). Due to communication through L2/3, the latency in L4 differed significantly between home and neighbor barrels with L4 stimulation (T = 3.4, p = 0.006) but not with L2/3 stimulation (T = 1.1, p = 0.32). L2/3 response latencies were not significantly different between home and neighbor barrels (L2/3 stimulation: T = 4.6, p = 0.0026; L4 stimulation: T = 1.5, p = 0.17), each constituting part of the midway point along the L4→L2/3→L4 pathway. Comparisons of velocity across stimulation sites and home versus neighbor barrels are included in Supplementary Results (Fig. S3A-B).

### Intra- and inter-barrel velocity

Response latencies increase with distance along pathways traced by axons. We used this relation to estimate conduction velocity. We constructed ROIs 18-30 μm wide within L4 to measure intralaminar conduction velocity within and between barrels (Fig. 7A: 30 µm wide ROIs). The heatmap in Fig. 7B shows responses to L2/3 stimulation that spread to L4 of the neighboring barrel. Plotting latency versus distance for the ROIs in L4, we find smooth propagation of responses to L2/3 stimulation (Fig. 7C). Fig. 7D displays traces from the home and neighbor barrels in Fig. 7A. For an example with L4 stimulation (sagittal, Fig. 7E: resting fluorescence; Fig. 7F: ΔF/F heatmap), we mapped latency in ROIs, color-coded as red-to-yellow, overlaid on the fluorescence image (Fig. 7G), demonstrating a gradient, particularly across the L4 neighbor barrel.

**Fig. 7:**
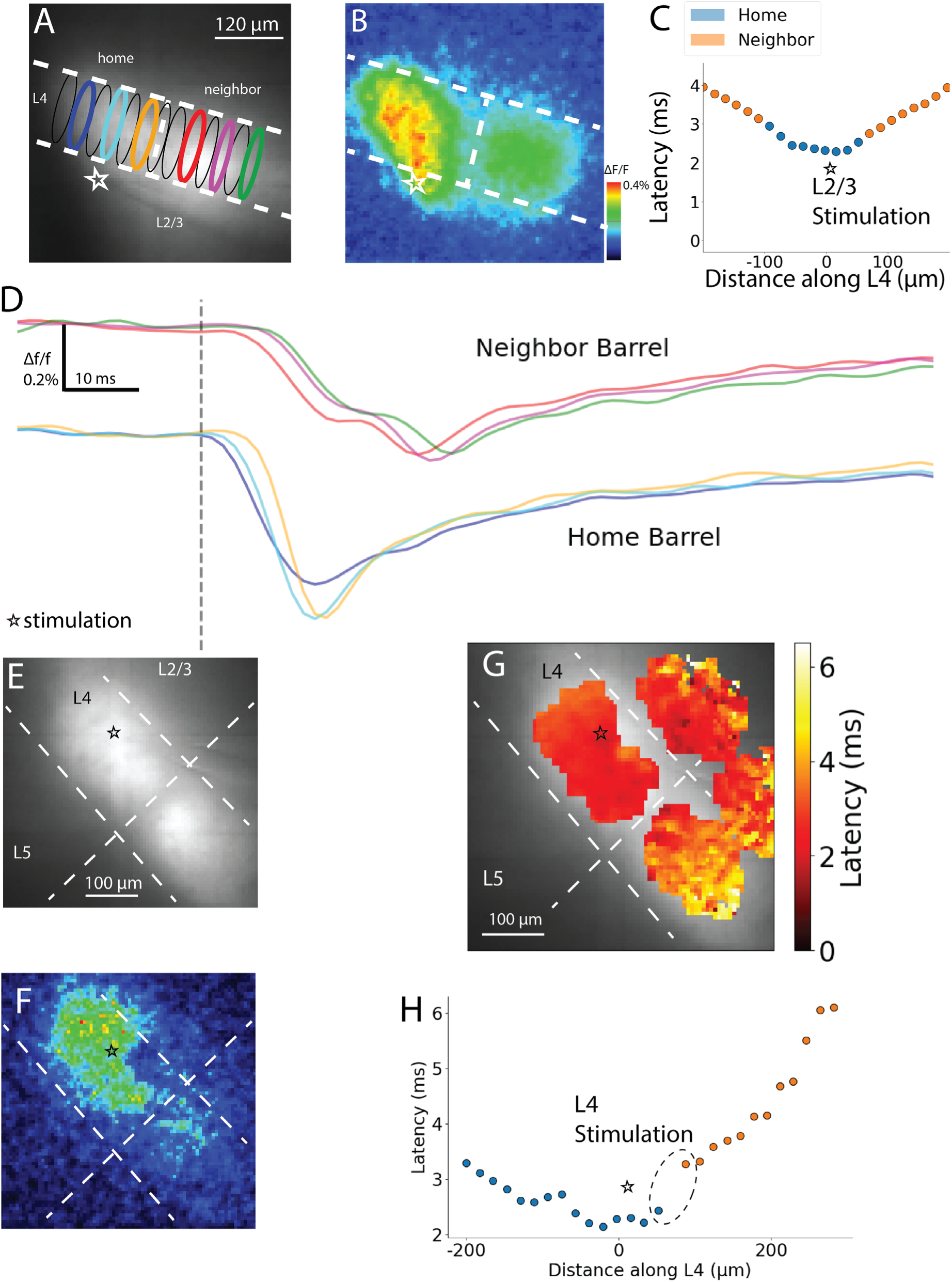
Velocity of propagation into neighbor barrels. **A**. Fluorescence image of a coronal slice showing ROIs used to compute velocity in home and neighbor barrels. **B**. Peak ΔF/F response map of the slice shown in A with stimulation site and layer/barrel boundaries annotated. **C.** Latency versus distance along L4 for responses to L2/3 stimulation, for a different slice from the one in A. Blue points are for the home barrel, while orange points are for neighbor barrels on either side. Domains of constant slope were used to determine velocity. **D**. ΔF/F traces from home and neighbor barrel ROIs of the corresponding colors in A arranged along the direction of propagation. **E.** Fluorescence image of a sagittal slice overlaid with the latency map, with the L4 stimulation site marked with a star. **F.** Peak ΔF/F response map with stimulation site and layer/barrel boundaries annotated for the slice shown in E. **G.** Heatmap of latency for the slice in E, laid over the resting fluorescence image. Latencies are shown only within the barrel ROIs, where fluorescence and thus responses are concentrated. **H**. ROIs from L4 in the example in E-G were used to plot latency versus distance along L4. The home and neighbor barrels show discontinuities in latency at the septum (black dotted ellipse).

In Fig. 7H, latencies along L4 (orange) show a jump of 0.60 ms over an 18-μm distance at the barrel boundary (black dotted ellipse), compared to the mean increment of 0.21 ± 0.04 ms per 18-μm in the rest of this plot. Such jumps were not seen for responses to L2/3 stimulation (Fig. 7C). However, other slices showed accelerations rather than delays in their L4 trajectories, or delays occurring within the neighboring barrel. These qualitative boundary changes in velocity mark discrete transitions from direct activation of the home barrel to indirect activation of the neighboring barrel, reflecting parallel paths of inter-barrel communication, including mono-or poly-synaptic signaling through L2/3 (Fig. 7C).

### Inter-barrel communication biases

To assess directional biases we compared inter-barrel relations in coronal (Fig. 1A-B) versus sagittal (Fig. 1C-D) slices. Successive whisker contacts occur several milliseconds apart, opposite to the direction of whisking. In particular, sagittal slices track barrels that map onto whiskers along the rostro-ventral direction toward which foveal protraction occurs, while barrels in coronal slices map onto rostro-dorsal whisking relevant to exploratory protraction. We determined how inter-barrel signaling varies in direction or phase (see Supplementary Results for slice plane variation). Fig. 8 compares barrel-averaged latency, half-width, neighbor/home ΔF/F ratio, and within-barrel conduction velocity across four directions (combining R-C and C-C). We also compared protraction-related directions (ventral: exploratory protraction; caudal: foveal protraction) to retraction-related directions (dorsal: exploratory retraction; rostral: foveal retraction) using the sum contrast model described in Supplementary Methods.

**Fig. 8.**
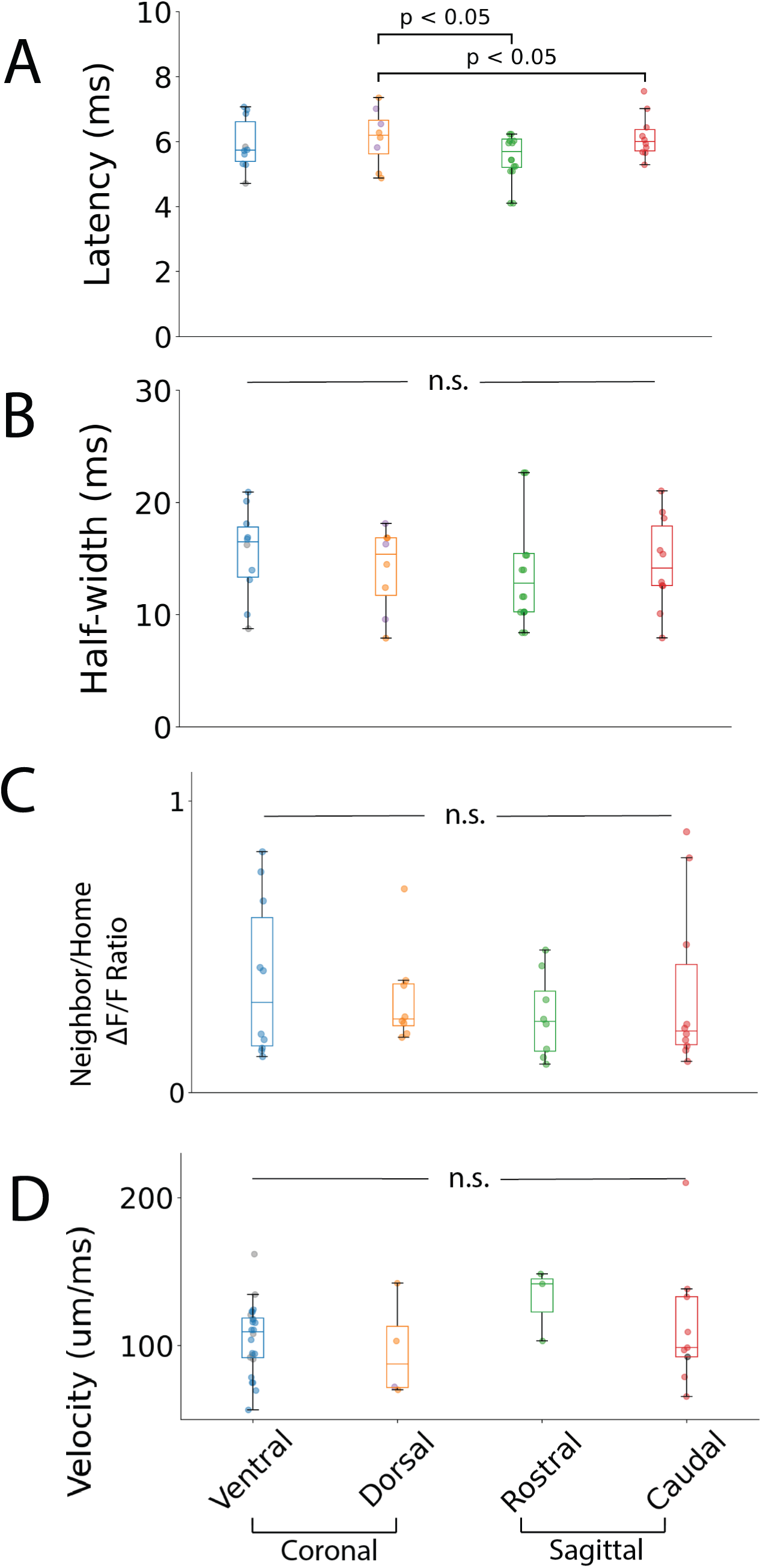
Comparison of responses between directions in barrel field. Latency (**A**), half-width (**B**), neighbor/home ΔF/F ratio (**C**) and velocity (**D**) of neighbor barrel responses grouped by propagation direction: rostral, caudal, dorsal, or ventral relative to the home barrel. Colors denote Caudal in M-S slices, Rostral in M-S slices, Dorsal in C-C slices, Ventral in C-C slices, Dorsal in R-C slices, and Ventral in R-C slices; R-C and C-C of the same direction are grouped in boxplots and scatter plots. Boxplots show minimum, first quartile, median, third quartile, and maximum with scatter; means ± SEM reported in Results. Statistical analysis used one-way ANOVA and Tukey’s HSD as appropriate.

Response latencies are consistent with the time between successive whisker contacts, and varied across directions (caudal: 6.2 ± 0.2 ms; rostral: 5.8 ± 0.4 ms; dorsal: 6.1 ± 0.3 ms; ventral: 5.9 ± 0.3 ms; mean effect size for contrast model: 0.23). These differences were significant (Fig. 8A; ANOVA, F = 5.9, p = 0.001, n = 10, 8, 8, 10 slices), with caudal > dorsal (Tukey’s HSD: p = 0.001) and dorsal > rostral (p = 0.001). Mean half-widths of neighbor barrel responses (Fig. 8B; caudal: 14.6 ± 1.3 ms; rostral: 13.5 ± 1.6 ms; dorsal: 14.0 ± 1.3 ms; ventral: 15.5 ± 1.3 ms; mean effect size: 0.65) were indistinguishable across directions (ANOVA, F = 0.5, p = 0.77, n = 10, 8, 8, 10, respectively). We normalized each neighbor barrel ΔF/F to its home barrel ΔF/F (Fig. 8C; caudal: 0.35 ± 0.09; rostral: 0.26 ± 0.05; dorsal: 0.32 ± 0.06; ventral: 0.39 ± 0.09; mean effect size (contrast model): 0.24) and found no anisotropy in the ΔF/F ratios (ANOVA, F = 0.45, p = 0.72).

Fig. 8D compares propagation velocity within L4 of neighboring barrels in response to L4 stimulation. Direction did not significantly impact velocity (ANOVA, F = 0.69, p = 0.56; ventral: 104 ± 5 µm/ms, n = 19; caudal: 113 ± 14 µm/ms, n = 9, dorsal: 97 ± 17 µm/ms, n = 4; rostral: 131 ± 14 µm/ms, n = 3; mean effect size of contrast model: 1.25). While deflection-evoked signals travel faster along rows than arcs (Petersen et al., 2003), and whisking velocity is greater along the rostral-caudal axis (Petersen et al., 2020), our velocities were more relevant to local circuitry. For instance, traversing wider barrels in the caudal aspect of BC (Fig. S1D-E) may require faster conduction. Conduction velocity, half-width, and ΔF/F of responses to single-pulse stimulation appear isotropic across BC, while variations in inter-barrel latency were significant.

### Paired-pulse depression

Whisking elicits repetitive activity at characteristic frequencies, and short-term synaptic plasticity contributes to processing these inputs. We investigated short-term plasticity of L4→L4 communication by stimulating L4 with paired pulses and evaluating the response ratio (PPR) in L4 of neighboring barrels. Temporal summation was evident for inter-pulse intervals less than 80 ms (Fig. 9A) so we subtracted single-pulse controls from the paired-pulse responses (Fig. 9B: dark blue traces). The subtracted traces reveal paired-pulse depression and a slow, incomplete recovery over IPIs from 20-120 ms. Figs. 9C-D plot PPR versus IPI for neighboring barrel responses in the four different barrel field directions. IPI significantly affected PPR (2-way ANOVA, PPR ∼ IPI + C(Direction) + IPI:C(Direction), F = 5.3, p = 0.021), and direction had a significant impact (F = 3.2, p = 0.012). We fitted these PPR versus IPI plots to the function 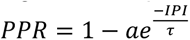 to determine the recovery time constant, τ and intercept at IPI = 0 ms. The qualities of fits were modest for dorsal (R² = 0.20), ventral (R² = 0.14), and rostral (R² = 0.28), but poor for caudal (R² = 0.002).

**Fig. 9:**
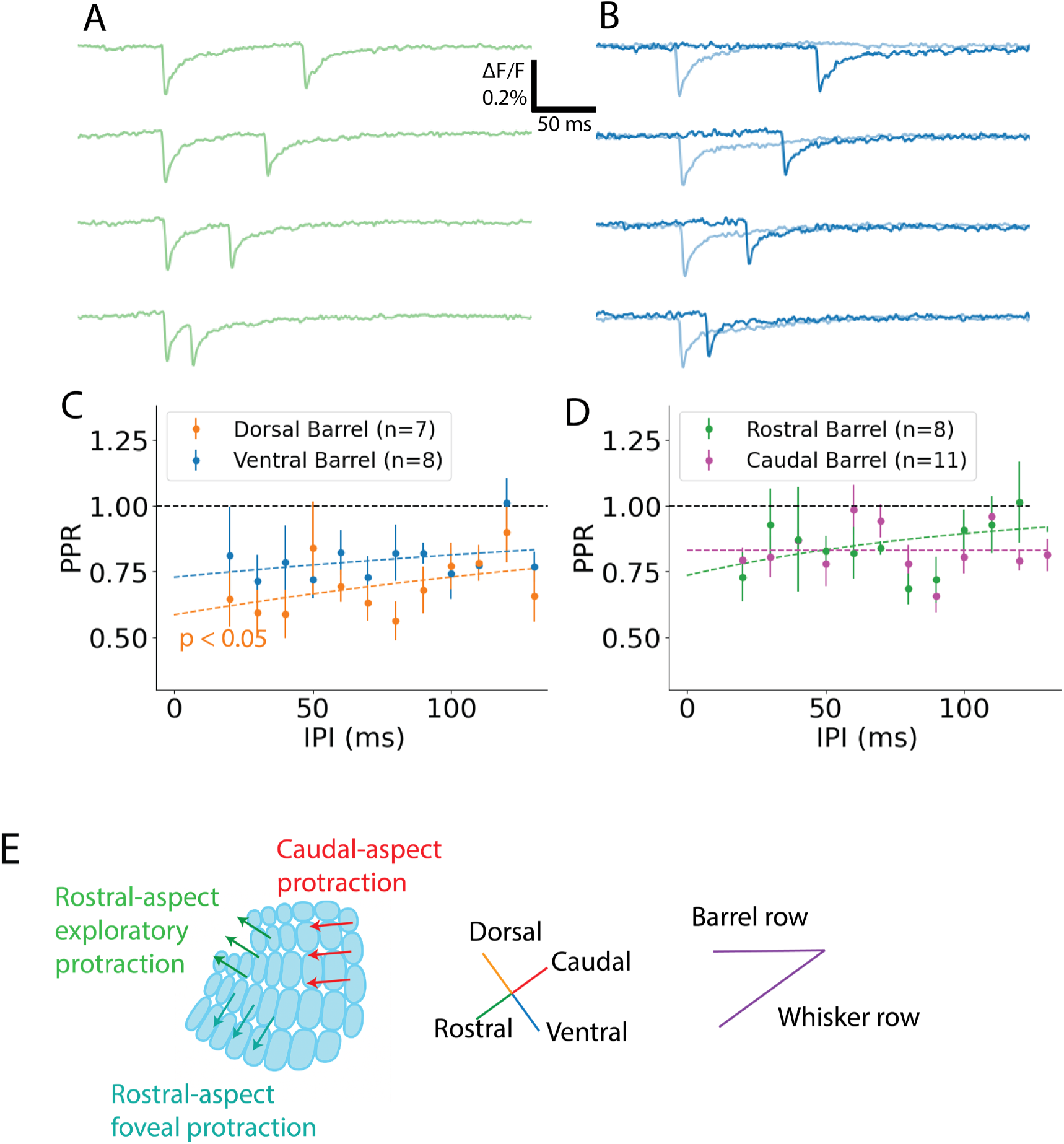
Paired-pulse ratio in neighbor barrels across directions. **A** Fluorescence traces from barrel delimited ROIs in L4 for paired-pulse stimulation at four different inter-pulse intervals (IPI). At sufficiently small IPIs, the first response does not return to baseline before the second pulse, leading to temporal summation that increases the second response. **B**. Single-pulse control traces are recorded (light blue traces) immediately before or after (determined randomly) the paired-pulse traces. Subtracted traces (dark blue), computed as paired-pulse minus single-pulse, remove temporal summation. **C-D.** Paired-pulse ratio (PPR, the amplitude of the second response divided by the first) plotted as a function of IPI for each direction, for coronal (**C**) and sagittal (**D**) slices. Solid curves represent exponential fits to recovery (see text). **E**. Diagram relating the kinematics of whisking patterns to barrel layout, showing that the dorsal-ventral slice plane in coronal slices tends to align along exploratory whisking, and the rostral-caudal slice plane in sagittal slices aligns along foveal whisking (Knutsen et al., 2005; Petersen et al., 2020). Orientation of four directions is based on the Brain Globe 3D atlas of barrel cortex viewed in coronal and sagittal planes (Claudi et al., 2020).

Table 1 summarizes the intercepts and recovery time constants. The mean effect size among these groups is 0.84 (contrast model). All four directions show significant depression, with the PPR intercept significantly less than 1 (Dorsal: T = 5.6, p = 0.0006, ventral: T = 5.4, p = 0.0009, caudal: T = 3.2, p = 0.005, rostral: T = 3.1, p = 0.009). The intercept varies significantly (ANOVA, F = 8.7, p = 0.0003), and is lower in the dorsal direction (Tukey’s HSD: caudal > dorsal, p = 0.001; rostral > dorsal, p = 0.032; ventral > dorsal, p = 0.025). Notably, the protraction PPR intercept was significantly greater than that of retraction (sum contrast-coded linear regression one-sided t-test, T = 6.3, p(30) < 0.001). Specifically, ventral PPR is greater than dorsal in foveal whisking, and caudal greater than rostral in exploratory whisking, yielding a consistently greater protraction PPR. During the protraction phase of free whisking this weaker paired-pulse depression in the caudal direction may preserve temporal fidelity for information moving along this specific direction.

**Table 1.** The PPR intercept at 0 ms and the exponential recovery time constant τ for each direction of inter-barrel propagation, determined by fitting the equation in the text to the plots in Fig. 9. The caudal direction did not show measurable recovery within the 120 ms IPI testing range, so τ was not determined. Values presented as mean ± SEM (n = number of slices).

| | PPR intercept, 1-a | recovery time constant $\tau$ (ms) | n |
| --- | --- | --- | --- |
| Dorsal | $0.58 \pm 0.08$ | $235 \pm 150$ | 7 |
| Ventral | $0.73 \pm 0.06$ | $269 \pm 230$ | 8 |
| Caudal | $0.83 \pm 0.06$ | ND | 8 |
| Rostral | $0.73 \pm 0.12$ | $110 \pm 85$ | 11 |

Direction also significantly impacted τ (ANOVA, F = 8.2, p = 0.002). Although dorsal and ventral barrel τ values were indistinguishable (Tukey’s HSD: p = 0.9), they were twice the rostral value (rostral versus ventral, p = 0.053; dorsal versus rostral, p = 0.15). In contrast, as suggested by the low quality of the exponential fit, caudal barrels do not show recovery within this range of IPIs, and show the least depression among the various directions (Fig. 9C). The illustration in Fig. 9E relates the directions between barrels to the angular offset of the whisking field, showing the whisking phase relevant to each BC slice plane. This illustrates the relevance of caudal and ventral directions to foveal and exploratory protraction, respectively, and of rostral and dorsal directions to foveal and exploratory retraction, respectively. The distinction between foveal and exploratory is most relevant to the fully protracted whisker setpoint, whereas whisker elevation is more constant near the caudal setpoint (Knutsen et al., 2005).

## Discussion

We imaged evoked voltage changes in Scnn1a-expressing excitatory neurons in mouse BC slices to characterize inter-barrel communication. Varying the stimulation site revealed that neurons in L2/3 relayed L4 excitation between adjacent barrels. Inter-barrel latency was longest in the caudal direction, but neighbor barrel conduction velocity, half-width, and amplitude ratio were isotropic. Paired pulse depression was asymmetric, with greater depression in the retraction direction. These findings reveal direction-, frequency-, cell type- and layer-specific mechanisms shaping somatotopic information flow in the BC, advancing our understanding of cortical sensory processing.

### Mechanistic insights into inter-barrel signaling

L4 stimulation elicited AMPA receptor-dependent responses in L4 of neighboring barrels. In contrast, L2/3 stimulation elicited AMPA receptor-independent responses in L4 of both home and neighboring barrels. This pattern is consistent with antidromic activation of L4 pyramidal neurons via the extensive L2/3 inter-barrel arborization (Feldmeyer et al., 2002; Staiger & Petersen, 2021) and previously-noted functional independence of barrels in L4 (Hill & Greenfield, 2014; Laaris & Keller, 2002; Petersen & Sakmann, 2001; Sato et al., 2008). Multiple inter-barrel pathways between L4 are likely, e.g. 𝐿𝐿4 → 𝐿𝐿2/3^ℎ𝑜*me*^ → 𝐿𝐿2/3*^neighbor^* → 𝐿𝐿4 or 𝐿𝐿4 → 𝐿𝐿2/3^ℎ𝑜me^ → 𝐿𝐿4; with lesser contributions from 𝐿𝐿4 → 𝐿𝐿2/3*^neighbor^* → 𝐿𝐿4. L5 may also contribute, but we focused on L4 and L2/3, and did not study L5. NBQX not only blocked neighboring barrel responses to L4 stimulation but also narrowed response half-widths, resolving synaptic response components. NBQX also revealed an inhibitory action: increased response amplitudes suggested that AMPA receptor-mediated activation of interneurons engages feedforward inhibition within and between barrels.

### Comparison to previous intracortical velocity studies

Our velocities (104-131 μm/ms) were generally faster than previous *in vivo* measurements ranging from 33-95 μm/ms (Armstrong-James et al., 1992; Petersen et al., 2003; Reyes-Puerta et al., 2016; Vilarchao et al., 2018), although 200 μm/ms has been reported (Lippert et al., 2007). Conduction velocities in BC slices tend to be greater: 74-473 μm/ms (Scheuer et al., 2023), 160 μm/ms (Haupt, 2000), and 130 μm/ms (Laaris & Keller, 2002), likely because naturalistic stimuli recruit cortical neurons more gradually (Ma & Patel, 2021), while electrical stimulation induces abrupt, synchronous depolarization. While natural responses reflect system-level function and longer-range connections, electrical stimulation in vitro reveals the underlying intracortical axonal conduction velocities and synaptic delays of microcircuit components. Repetitive stimulation (100-500 Hz) *in vivo* leaves velocities unaffected (113 μm/ms (Ferezou et al., 2006; Margalit & Slovin, 2018), consistent with constant velocities between successive pairs of pulses (Fig. S3D).

### Directional anisotropy and propagation dynamics

Our experiments probed barrel response properties for directional preference. These preferences may be relevant to differences between fast foveal (with rostro-ventral protraction) versus slow exploratory (with rostro-dorsal protraction) behavior. All neighbor barrel single-pulse response properties except latency were isotropic. Caudal latencies were longest. Whisker contact with an object is communicated to its caudal neighbor barrel approximately 10 ms after the adjacent caudal whisker contacts the same object (Vilarchao et al., 2018). This asymmetry parallels the lower density of barrel projections from L4 neurons to L2/3 caudal neighbors (Bernardo et al., 1990); as the number of axons grows, the minimum axonal path length decreases, reducing latency.

Caudal latencies relate to inter-barrel integration during protraction, and dorsal latencies relate to integration during retraction. Long, caudal whiskers may specialize in slow, exploratory whisking compared to the small, rostral whiskers preferentially recruited during foveal whisking (Hobbs et al., 2016). However, whisker specialization is manifold. Caudal barrels (e.g. in M-S slices) sense the animal’s periphery, responding faster to strong, unexpected peripheral stimuli.

Barrel size may also be relevant, with larger barrels requiring faster conduction to function as a unit. Rostral barrels were 17% smaller than caudal barrels (Fig. S1D-E). Larger barrels, corresponding to the longest and thickest whiskers, receive stronger excitation (Grant et al., 2009; Woolsey, 1970). Since speed increases with signal amplitude across cortical areas (Afrashteh et al., 2021), these highly-excited larger barrels may require faster barrel-wide propagation.

### Anisotropic short-term depression

Paired-pulse stimulation revealed significant direction-dependent variations in inter-barrel short-term synaptic plasticity, and a modest amount of this variance was captured in a simple single-exponential model. Retraction-related responses showed the strongest depression at short IPIs; protraction-related responses, especially caudal, showed the least depression. Protraction is primarily directed rostrally, but also may extend ventrally near the rostral-most setpoint for whisking against a vertical pole (Petersen et al., 2020). This implies foveal protraction, fixated on an object, and involving successive dorsal contacts. In contrast, exploratory protraction in free air produces a dorsal rather than ventral change in whisker elevation (Knutsen et al., 2005). The higher PPR in protraction—especially caudal—directions will enhance transmission of repetitive activity in these directions.

IPIs of 20-100 ms are relevant to repetitive thalamocortical input from the same whisker, falling into the 10-50 Hz unevenly-distributed range that is sub-linearly boosted during thalamic processing in the 5-25 Hz whisking range (Sosnik et al., 2001). Repetitive L4 input propagates to neighboring barrels, during which inter-barrel depression of communication becomes relevant. This communication is attenuated more strongly in the retraction-related direction, preserving the fidelity of protraction phase transmission. This asymmetry enhances multi-barrel integration during information-rich protraction which lasts 30% longer than retraction (Bermejo et al., 2002; Leiser & Moxon, 2007). The shorter retraction phase increases the synchrony of excitatory input, which requires stronger attenuation in preparation for the next protraction. This directionally-biased excitatory circuit can thus exploit unequal phase durations to enhance sensory processing.

## Conclusion

Combining genetically-targeted voltage imaging with a directional alignment of BC-whisker anatomy connected population-level responses to system-level sensory processing. Inter-barrel communication between L4 excitatory neurons employs layer-specific pathways that depend on direction and frequency. The L4→L2/3→L4 circuit supports inter-barrel signaling, and its anisotropy in latency and in synaptic depression tune intracortical communication to the spatiotemporal structure of whisker input. These mechanisms serve in the interpretation of whisking phases, patterns, and frequency. More broadly, this study suggests that spatiotemporal alignment to sensory input may be relevant to how microcircuits implement cortical maps. Extending this investigation to the single-cell level with sparse CreERT2-driven labeling (Feil et al., 2009) and single-cell stimulation (Canales et al., 2022) have the potential to reveal the detailed circuitry underlying the processing of sensory inputs.

## Supporting information

Animation L4 stimulation

Animation L2/3 stimulation

## Acknowledgements

Anna White collected data for several NBQX experiments with the support of WISCIENCE/Cellular and Molecular Biology of Stress Summer Research Program and under the mentorship of the authors.

## Funding

Supported by NIH grant R35 NS127219, NIH training grant T32 GM130550.

Multimedia 1: Spatiotemporal spread of excitation across barrels and layers for L4 stimulation. Left: Animated sequence of heat maps of peak responses from just before stimulation to 10 ms post-stimulation, paralleling the heatmap time series of Fig. 6G for a separate example. The resting fluorescence image is shown at the animation end. Stimulation site is marked with a star. Right: Traces in the home (L4: blue, L2/3 green) and neighbor (L4: orange, L2/3: red) barrels. ROI outlines of the same color in maps correspond to traces. Dotted blue line marks stimulation time. With L4 stimulation, the response in L4 of the home barrel precedes that in L2/3 and in neighbor barrels.

Multimedia 2: Stimulation in L2/3 shows near-simultaneous activation of three unstimulated ROIs. Left: Heat maps of peak responses animated from just before stimulation to 10 ms post-stimulation, paralleling the heatmap time series of Fig. 6H for a different slice. The resting fluorescence image is shown at the animation end. Stimulation site is marked with a star. Right: Traces in the home (L4: blue, L2/3 green) and neighbor (L4: orange, L2/3: red) barrels. ROI outlines of the same color in maps corresponding to traces. Dotted blue line marks stimulation time. Biphasic responses reflect multiple L2/3-L4 pathways mediating inter-barrel communication.

## Supplementary information for

### Supplementary Methods

*Analysis, Statistics.* PhotoZ implements baseline subtraction, temporal and spatial filtering, and linearly-interpolated measurements of latency, half-width, and peak ΔF/F measurements. The latencies are times from the stimulation pulse to half-amplitude. Time windows for analysis were from 3 ms before stimulation to 50 ms post-stimulation, with independent samples obtained as barrel-averaged measurements. Domains with lengths of at least 50 μm showing constant velocity were identified for linear regression and velocity measurement. Velocity was calculated with the SciPy.stats Python library with linear regression on latency versus distance along L4, using 7 or more regions of interest (ROIs) in each barrel. Pixels within 50 μm of the stimulation site were excluded. Septa between barrels were also excluded due to their dim fluorescence.

We tested multiple groups for significance with one-way or two-way ANOVA (SciPy.stats). If ANOVA indicated significant variation, Shapiro-Wilk tests confirmed normality before we used Tukey’s honestly significance difference (HSD) test for pairwise comparisons (Statsmodels, Python). We confirmed that inter-class variance was not significantly greater than intra-class variance with a mixed-effect linear regression model implemented in R, obviating the need to control for pseudoreplication effects between slices of the same animal or litter (Lazic, 2010). In cases where only two groups were compared, we used one-sided Welch’s t-tests (Scipy.stats). We also used one-sample one-sided t-tests (Scipy.stats) to compare mean against null hypothesis values where relevant.

To test the dependence of measurements on direction we hypothesized that the differences between dorsal-minus-ventral and rostral-minus-caudal will share the same sign because ventral and caudal neighbor activations are associated with whisker protraction, whereas dorsal and rostral neighbor activations are associated with retraction, taking the angular offset between BC and whisker field into account. Because each group is comprised of independently measured barrels, we used a contrast coding system of linear regression (Patsy Python:sum contrast coding; Statsmodel’s ordinary least squares function). Using this sum contrast coding (comparing to the group mean as reference), we tested whether the differences of each pair of group means (represented by μ_i_) would sum to significantly greater than zero on average (one-sided t-test), i.e. alternative hypotheses of:

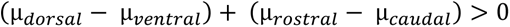

This tested for protraction < retraction; signs were flipped to test for protraction > retraction. The contrast model allowed for specific testing without the family-wise error rate corrections in Tukey’s HSD test.

We designed our sample sizes to detect intracortical processing differences on the order of at least 15%. To achieve a standard power level (0.8) and false positive rate of 5%, allowing for an uncertainty of 8%, we calculated that a minimum sample size of 8 is needed: 𝑛_2_ = 𝑛_1_ = 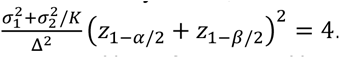. For post-hoc power analysis, we measured variances (ΔF/F ratios: 14%, half-width: 25%, velocity: 19-26%, latency: 10-19%, paired-pule ratio (PPR) parameters: 20-50%) and calculated effect sizes and sample sizes based on the contrast model (reported in Results), which are similar to those of the sagittal-coronal comparison. Tests of half-width (0.15), ΔF/F ratios (0.17), and latency (0.54) were underpowered. The power levels of velocity (1.0) and PPR at 20 ms interpulse interval (IPI) (0.73) indicated a sufficient sample size to achieve standard statistical power.

### Supplementary Results

Our supplementary results include further details on barrel-to-whisker alignment, an analysis of barrel dimensions of each type of slice, a summary of non-significant results for response properties grouped by direction, and results demonstrating controlled variables do not have unexpected effects on the velocity and PPR results.

As illustrated in Fig. 1, we categorized our slices by their preservation of connections. The connections are preserved depending not only on the plane of section but also on slice origin (e.g. coronal sections from rostral versus caudal aspects, as in Fig. 1A). This produces the four categories specified in Results: rostral-aspect coronal (R-C slice, Fig. 1A, 1B), caudal-aspect coronal (C-C slice, Fig. 1A, 1C), lateral-aspect sagittal (L-S slice, Fig. 1D, 1E), and medial-aspect sagittal (M-S slice, Fig. 1D, 1F). R-C slices tend to present a cross-section of rows C, D, or E (labeled in Fig. 1B, bottom right), which curve into the rostro-dorsal direction in the rostral aspect. (Fig. 1A, right). These rows have narrow septa and higher barrel count (7-10 per slice). In contrast, C-C slices present fewer, larger barrels and septa (Fig. 1C). Sagittal slices originated from either aspect (Fig. 1E: M-S, Fig. 1F: L-S), with lateral slices containing barrels corresponding to ventral whiskers and medial slices containing barrels corresponding to dorsal whiskers (Petersen et al., 2020). In L-S slices, however, the slice plane becomes nearly tangential, so these slices were excluded.

To evaluate the slice alignment illustrated in Fig. 1, we sampled the Brain Globe 3D atlas (Claudi et al., 2020) in 12 locations relevant to each group of slices to estimate the probability *P(row)* of adjacent barrels being in the same row. We obtained *P(row) =* 0.90 in R-C slices, 0.50 in C-C slices, and 0.58 in M-S slices, while P(*arc) =* 0.06, 0.40, 0.17 (note *P(row) + P(arc) + P(diagonal) = 1*). We originally intended to associate coronal slices with arcs and sagittal slices with rows. However, this atlas analysis suggests that our sections align well to foveal and exploratory whisking patterns. R-C and C-C slices map onto exploratory whisking, in which protraction is angled in the rostral-dorsal direction. Meanwhile, the sagittal plane can be used to track foveal whisking, in which protraction tends toward the rostral-ventral direction (Knutsen et al., 2005; Petersen et al., 2020). This correspondence is a consequence of the angular offset between barrel and whisking fields. Our alignment allows us to relate these whisker field axes to inter-barrel communication.

**Fig. S1:**
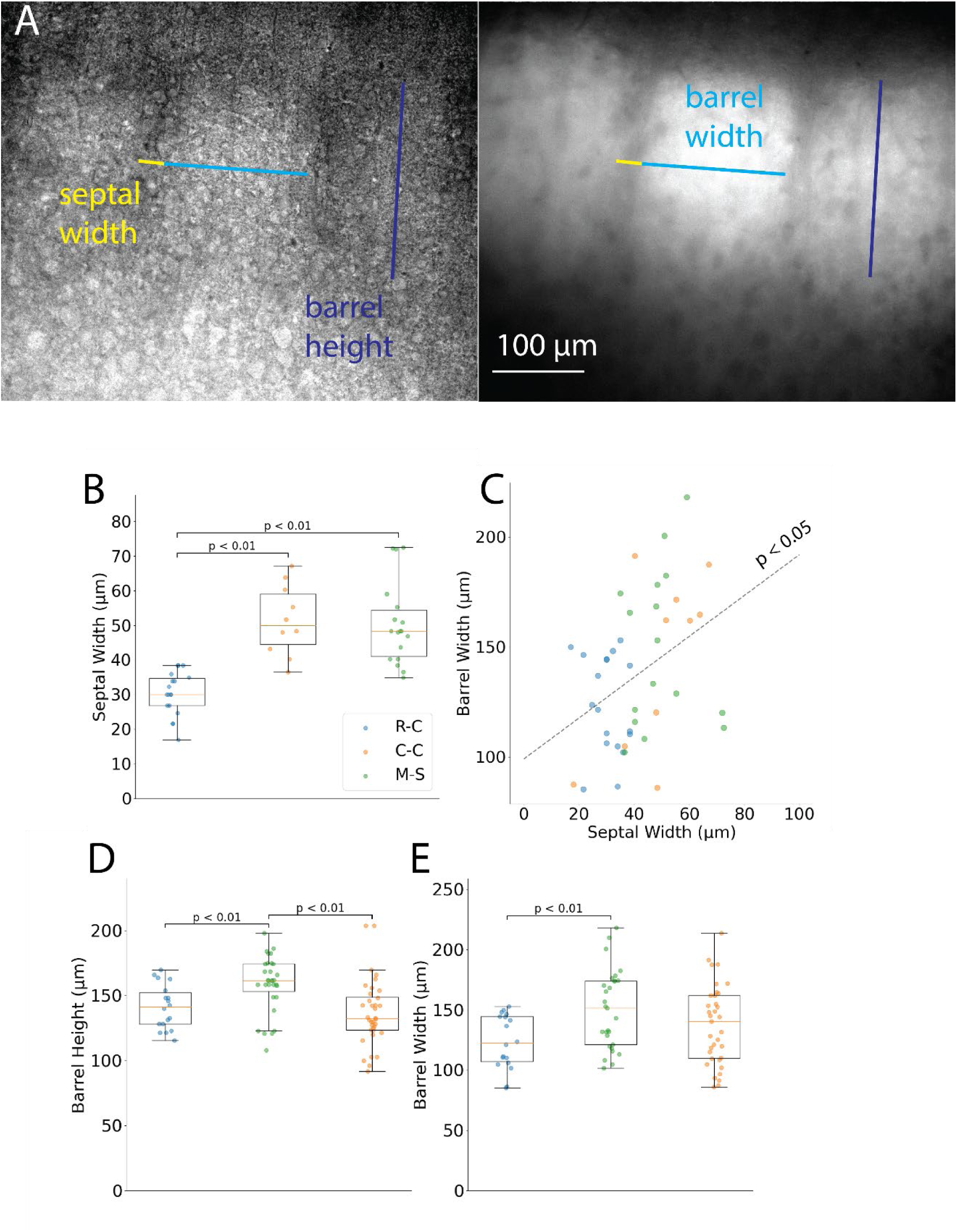
Barrel dimensions across slice types. **A.** Bright field (left) and resting fluorescence (right) images featuring barrel width (light blue) and height (dark blue), and septum width (yellow). The L2/3-L4 boundary is determined based on the presence of septal columns and higher fluorescence in L4 than in L2/3. **B**. Comparison of septal width among R-C, C-C, and M-S slices (note that L-S slices were not included, see text). **C**. Plot of barrel width versus septum width with line from linear regression (p < 0.05). **D-E.** Barrel height and width differ across slice types (M-S barrel height is greater than that of R-C and C-C; R-C barrels were narrower than M-S barrels). Each point indicates one barrel or septum; lines show mean ± SEM. Significance assessed by Welch’s t-test.

#### Barrel dimensions

The clear boundaries highlighted by Scnn1a labeling enabled us to estimate the basic barrel dimensions. The septal width, barrel width, and barrel height, annotated on bright-field (left) and fluorescence (right) images in Fig. S1A, were estimated from the cytoarchitectural boundaries. We distinguished L2/3 from L4 based on a steep fluorescence gradient and the loss of distinction between septa and barrel hollows at the boundary. Fig. S1B compares septal widths between the R-C, C-C, and M-S slices. The mean septal width in the rostral row of R-C slices, 30.3 ± 1.5 µm (n = 18 septa, 15 slices), was significantly smaller than in C-C slices (51.5 ± 3.2 µm, n = 10 septa, 10 slices; Welch’s t-test: T = 5.99, p = 5×10^−5^) or in M-S slices (50.4 ± 2.8 µm, n = 18 septa, 13 slices, T = 6.38, p = 10^−6^). R-C and M-S septal widths did not differ significantly (T = 0.24, p = 0.8). These variations were exploited to track the transition from rostral-aspect (R-C) to caudal-aspect (C-C) in the sequence in which coronal slices were cut. Barrel width and height did not correlate significantly (linear regression, R = 0.19, p = 0.09), which is expected given the heterogeneity of barrel dimensions and the fact that the thickness of L4 limits barrel height to 208 ± 5 µm or less (DeFelipe et al., 2002). However, barrel and septal width showed a significant positive correlation (Fig. S1C, R = 0.38, p = 0.01). Additionally, sagittal slices were cut at a smaller angle (less than a right angle) to the cortex layers, particularly towards the medial aspect but also in the lateral aspect. As a result, L4 is thicker in some sagittal slices and barrel height is significantly greater in M-S slices (Fig. S1D, 159 ± 4 µm, n = 31 barrels) than in R-C slices (141 ± 4 µm, n = 18, T = 3.1, p = 0.003) or in C-C (137 ± 4 µm, n = 36, T = 3.9, p = 0.0003). Barrel widths were significantly smaller in R-C slices (Fig. S1E, 124 ± 5 µm, n = 18) than in M-S slices (150 ± 6 µm, n = 29, T = 3.33, p = 0.002), but not significantly different from C-C slices (138 ± 6 µm, n = 35, T = 1.55, p = 0.13). These differences in barrel width match the trend in barrel size seen in the tangential view of BC, in which barrels become smaller in the rostro-medial aspect and have smaller cross-sections in the coronal plane (Fig. 1B, lower right; (Petersen et al., 2020). Our choice of 200 µm slice thickness limits background from barrels outside of the microscope focal plane, given the sum of our measurements of barrel width plus septal width (154-200 µm).

#### Spatiotemporal biases in inter-barrel communication: isotropic response properties

In Fig. S2A, comparisons of mean latency of neighbor barrel responses showed no significant differences between planes (Fig. 8E, Welch’s t-test, T = 1.7, p = 0.11) or between protraction and retraction phases (T = 0.63, p(32) = 0.27).

In Fig. S2B, the contrast model showed no significant difference in half-width between protraction and retraction directions (T = 1.0, p(32) = 0.16), nor between coronal and sagittal slices (Welch’s t-test, T = 0.75, p = 0.46).

Direction did not significantly affect neighbor/home ΔF/F ratio, nor did slice plane (Welch’s t-test, coronal: 0.36 ± 0.05; sagittal: 0.31 ± 0.05; T = 0.67, p = 0.51) or phase (Fig. S2C; sum contrast model, T = 1.0, p(32) = 0.15).

Grouped by slice plane, the smaller mean coronal velocity (117 ± 5 µm/ms, n = 29) was not significantly different from the mean sagittal velocity of 133 ± 14 µm/ms, n = 12 (Fig. S2D; T = 1.1, p = 0.30). With its more specific test, the contrast model showed that protraction velocities were not significantly smaller than retraction velocities (T = 0.12, p(36) = 0.45).

**Fig. S2:**
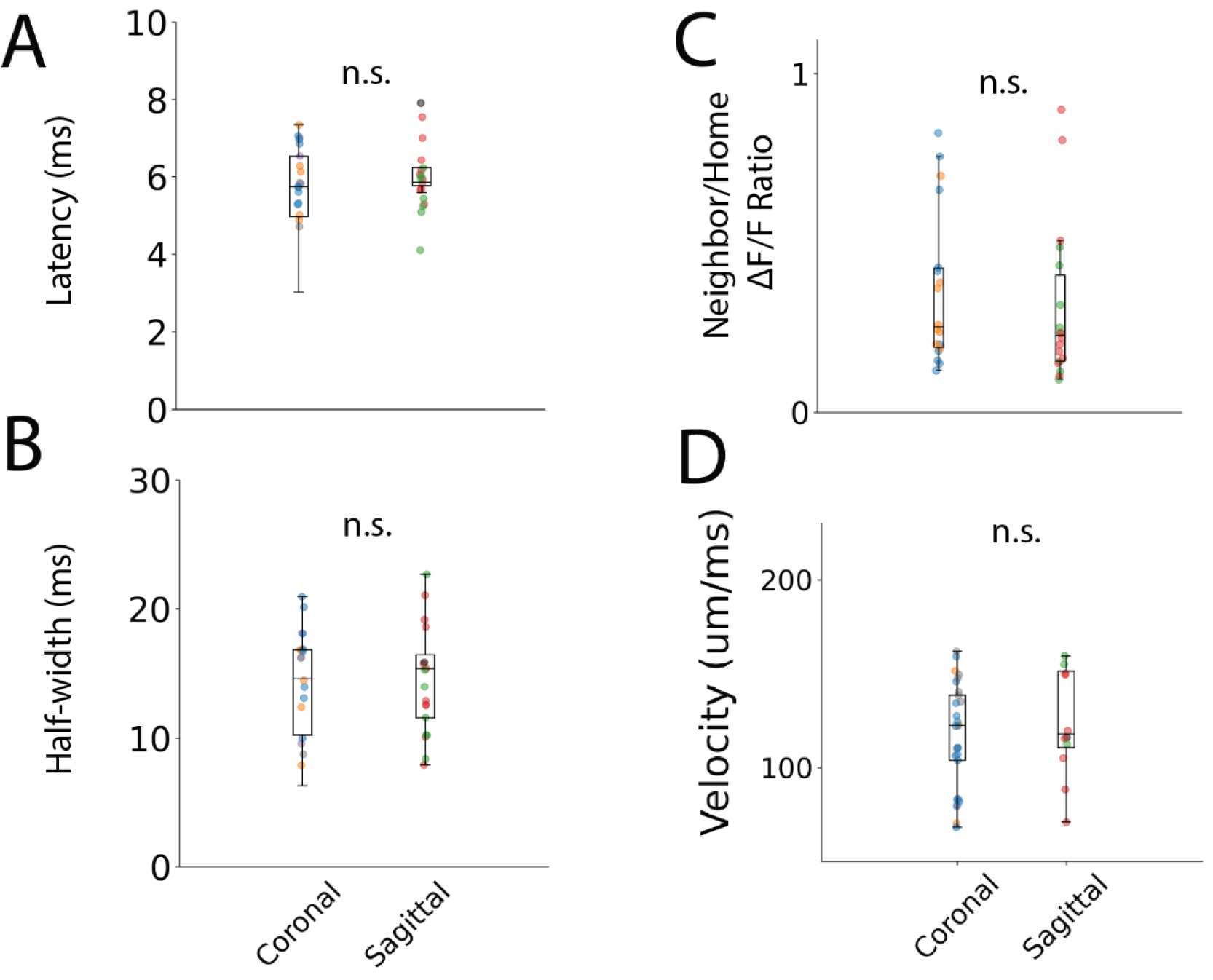
Inter-barrel response properties are symmetric in coronal and sagittal slices. Latency (**A**), half-width (**B**), neighbor/home ΔF/F ratio (**C**) and velocity (**D**) of neighbor barrel responses grouped by slice plane: coronal or sagittal. Colors denote Caudal in M-S slices, Rostral in M-S slices, Dorsal in C-C slices, Ventral in C-C slices, Dorsal in R-C slices, and Ventral in R-C slices; R-C and C-C of the same direction are grouped in boxplots and scatter plots. Boxplots show minimum, first quartile, median, third quartile, and maximum with scatter; means ± SEM reported in Results. Statistical analysis used Welch’s t-tests as appropriate.

#### Velocity is unaffected by AMPA-receptor blockade or synaptic depression

The mean velocity within L4 neighbor barrels was similar regardless of stimulation site (Fig. S3A, open circle markers; L2/3: 129 ± 12 µm/ms, L4: 122 ± 5 µm/ms, T = 0.98, p = 0.36). L4 neighbor velocities were computed for the minority of cases where NBQX did not completely abolish responses (ACSF: n = 31 responding L4 neighbor barrels, NBQX: n = 6). NBQX had no significant effect on velocity (Fig. S3A, bordered markers; ACSF and NBQX, L2/3: T = 1.16, p = 0.27, L4: T = 1.0, p = 0.34). This suggests that axonal conduction speed (antidromic or orthodromic) determines the velocities of response spread, despite the biphasic nature of many responses in neighboring barrels (Fig. 7D, top traces). Velocities were greater in the home barrel (132 ± 10 µm/ms) than the neighboring barrel with both L2/3 or L4 stimulation (mean: 123 ± 6 µm/ms), but this difference was not significant (T = 0.77, p = 0.45, Fig. S3B).

Depression in amplitude was not accompanied by a change in half-width (Fig. S3C, home: T = 0.14, p = 0.89; neighbor: T = 0.66, p = 0.51). Additionally, there was no change in conduction velocity from the first to second pulse (Fig. S3D, 99 ± 9 μm/ms to 98 ± 8 μm/ms, T = 0.11, p = 0.91), suggesting a constant speed for processing information between adjacent barrels.

**Fig. S3:**
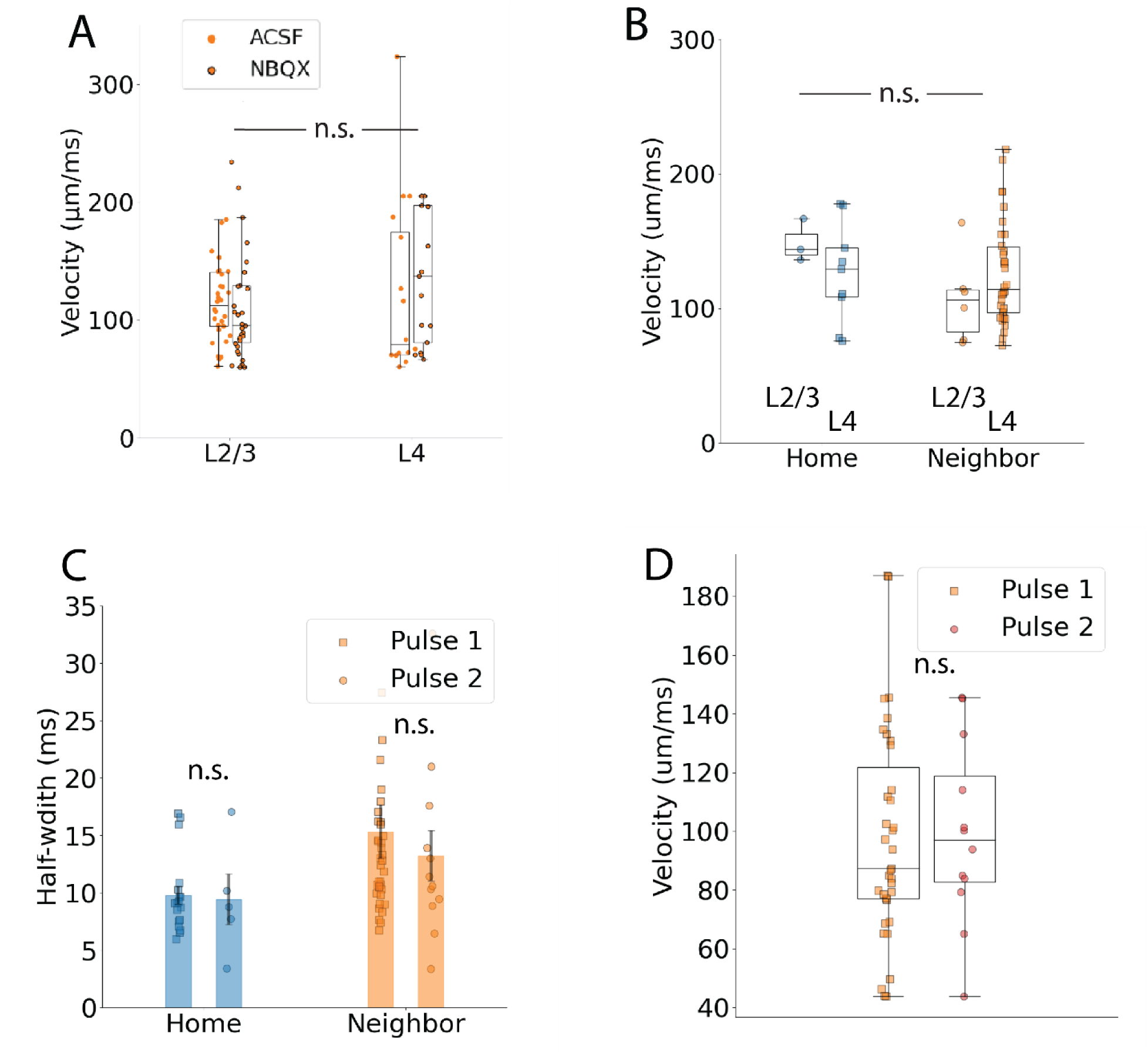
Velocity and half-width persist across manipulations. **A**. Boxplot and scatter of velocity measurements before and after NBQX. NBQX had no significant effect. **B.** Velocity was greater in the home barrel but not significantly different from neighbor barrel velocities. **C.** No significant change in velocity between the first and second responses. **D.** Response half-width was also unchanged. Note that the difference between neighbor and home half-widths was discussed in regard to Fig. 3H, so we have left it unmarked here to avoid repetition of results and to focus on the present comparison of first and second responses. Boxplots show minimum, first quartile, median, third quartile, and maximum with scatter; means ± SEM reported in Results. Statistics used barrel means as independent samples, although individual ROI measurements are displayed.

#### Age and sex do not affect response properties of the present study

A comparison of male versus female mice showed no significant differences in latency, half-width, velocity (Welch’s t-test, T < 0.001, p = 1.0 for all), or paired-pulse ratio (PPR) (multivariate linear regression, F=0.55, p =0.46). Mice aged 30 to 60 days showed no significant correlation between these metrics and age: response latency, half-width (linear regression, L2/3: latency: p = 0.23, half-width: p = 0.48; L4: latency: p = 0.16, half-width: p = 0.57), velocity (linear regression, p = 0.66) or PPR (multivariate linear regression, F = 0.006, p = 1.0).

